# O-GlcNAcase promotes dendritic spine morphogenesis while downregulating their GluA2-containing AMPA receptors

**DOI:** 10.1101/2025.08.15.670533

**Authors:** Linkun Han, Sabrina Galizia, Jingyu Pan, Olof Lagerlöf

## Abstract

Dendritic spines are essential for synaptic transmission, neural circuit organization, and cognitive function. Their morphology and density influence synaptic plasticity, learning, and memory. Many proteins in dendritic spines are modified with O-GlcNAc, a monosaccharide that can be attached and removed from serines and threonines. O-GlcNAc has been implicated in multiple brain disorders, yet the role of O-GlcNAcase (OGA), the enzyme that removes O-GlcNAc modification from proteins, in dendritic spine regulation remains unclear. This study examines the role of OGA in spine and synapse morphogenesis. Immunohistochemical and biochemical analyses reveal OGA present in dendritic spines. Functional assays show that OGA promotes spine maturation, increases spine density, and alters synapse size. Additionally, OGA modulates the α-amino-3-hydroxy-5-methyl-4-isoxazolepropionic acid receptor (AMPAR), down-regulating GluA2-containing receptors in developing and mature neurons. These findings highlight OGA as a key regulator of excitatory synaptic remodeling and a therapeutic target for synapse-related pathologies such as Alzheimer’s disease and autism.

## Introduction

Learning, memory, and overall cognitive function depends on synaptic plasticity, the ability of synapses to strengthen or weaken over time in response to neural activity[1]. Excitatory synapses are primarily located on dendritic spines. Dendritic spines are small protrusions on neuronal dendrites[2]. Spine structure and density are dynamically regulated and play a crucial role in synaptic function and plasticity. Changes in dendritic spine morphology are associated with both normal brain development and pathological conditions, including neurodevelopmental disorders such as autism spectrum disorder (ASD), schizophrenia, and neurodegenerative diseases like Alzheimer’s disease (AD) [3-5]. Understanding the molecular mechanisms governing spine morphogenesis is essential for elucidating the pathophysiology of these disorders and identifying potential therapeutic targets.

One important post-translational modification (PTM) involved in synaptic regulation is O-linked β-N-acetylglucosaminylation (O-GlcNAcylation), a dynamic and reversible modification in which a single N-acetylglucosamine (GlcNAc) moiety is added to serine or threonine residues of nuclear and cytoplasmic proteins[6]. O-GlcNAcylation is distinct from other glycosylation forms as it primarily is not involved in protein secretion but rather serves as a critical regulatory mechanism for intracellular signaling, metabolism, and gene expression[7-9]. The balance between O-GlcNAc addition and removal is maintained by two key enzymes: O-GlcNAc transferase (OGT), which catalyzes the addition of O-GlcNAc, and O-GlcNAcase (OGA), which removes the modification [6, 10]. OGT, OGA and O-GlcNAc are highly prevalent in neurons where many proteins associated with synaptic plasticity are modified, e.g. SynGAP and aCaMKII [11, 12]. Emerging evidence mainly based on pharmacological and genetic modulation of OGT suggests that O-GlcNAc modifies these proteins to control synaptic plasticity by affecting the synaptic abundance of AMPA receptors[13-15]. The AMPA receptor is an excitatory neurotransmitter receptor. The majority of AMPA receptors are tetramers formed by dimers of either the GluA1 and GluA2 subunits or GluA2 and GluA3 subunits[16, 17]. AMPARs mediate the majority of excitatory neurotransmission in the brain, and their regulation is crucial for synaptic strength and function[16]. Multiple studies have demonstrated that synapse size is positively correlated with the number of AMPARs, and that neuronal activity can modulate both synapse size and AMPAR content[18, 19]. Moreover, synapse size is another important determinant of synaptic strength. Larger synapses with increased postsynaptic density (PSD) components are generally associated with greater synaptic efficacy, whereas smaller synapses may undergo structural plasticity or pruning[20, 21]. In addition to its effect on AMPA receptors, genetic deletion of OGT has been shown to decrease synapse number without affecting synapse size[14]. Indeed, dysregulation of OGT has been linked to memory impairment, neurodevelopmental disorders and neurodegenerative diseases, highlighting the importance of O-GlcNAc in CNS function[22].

While OGT has been studied in the context of synaptic function, OGT has proven difficult to target pharmacologically in vivo. Several clinical trials are instead targeting OGA. However, the role of OGA in synaptic plasticity remains less well characterized. Previous studies using OGA knockout models have provided valuable insights—global OGA deletion in mice results in perinatal lethality, while neuron-specific knockouts have shown alterations in tau phosphorylation and neurodegeneration[23, 24]. Despite this, the subcellular localization and specific functional contributions of OGA within synapses have not been elucidated. Our study aims to address these gaps by specifically investigating OGA’s subcellular localization and function in excitatory synapse plasticity.

## Results

### OGA is present in dendritic spines

O-GlcNAc is highly expressed in neuronal synapses in the brain[25], and biochemical fractionation and immunohistochemistry studies have demonstrated the presence of OGT in neuronal synapses[14, 26, 27]. However, the precise localization of OGA in neurons remains largely unclear. To investigate the distribution of OGA in neurons, we performed immunohistochemical analysis of OGA on mouse brain sections of the hippocampus where the nucleus, soma and dendrites can be distinguished. We observed that OGA is localized within the dendritic compartment (Fig. 1A). To further examine its subcellular localization, we isolated biochemically different fractions from mouse brain hippocampus and conducted western blot analysis. Our results revealed that OGA is present in most subcellular fractions. However, in contrast to OGT, it is absent from the PSD and present at a lower level in synaptosomes compared to the whole-cell fraction (Fig. 1B). To gain additional insights into OGA localization, we transfected hippocampal neurons at DIV14 with expression vectors encoding GFP and subsequently immunostained the transfected neurons for OGA at DIV16. Interestingly, we observed OGA expression in specific regions of the dendrites (Fig. 1C), with a higher intensity in mature spines compared to immature spines (Fig. 1D). Collectively, these findings indicate that OGA is highly expressed in dendrites and in dendritic spines but not in the PSD.

**Figure 1.**
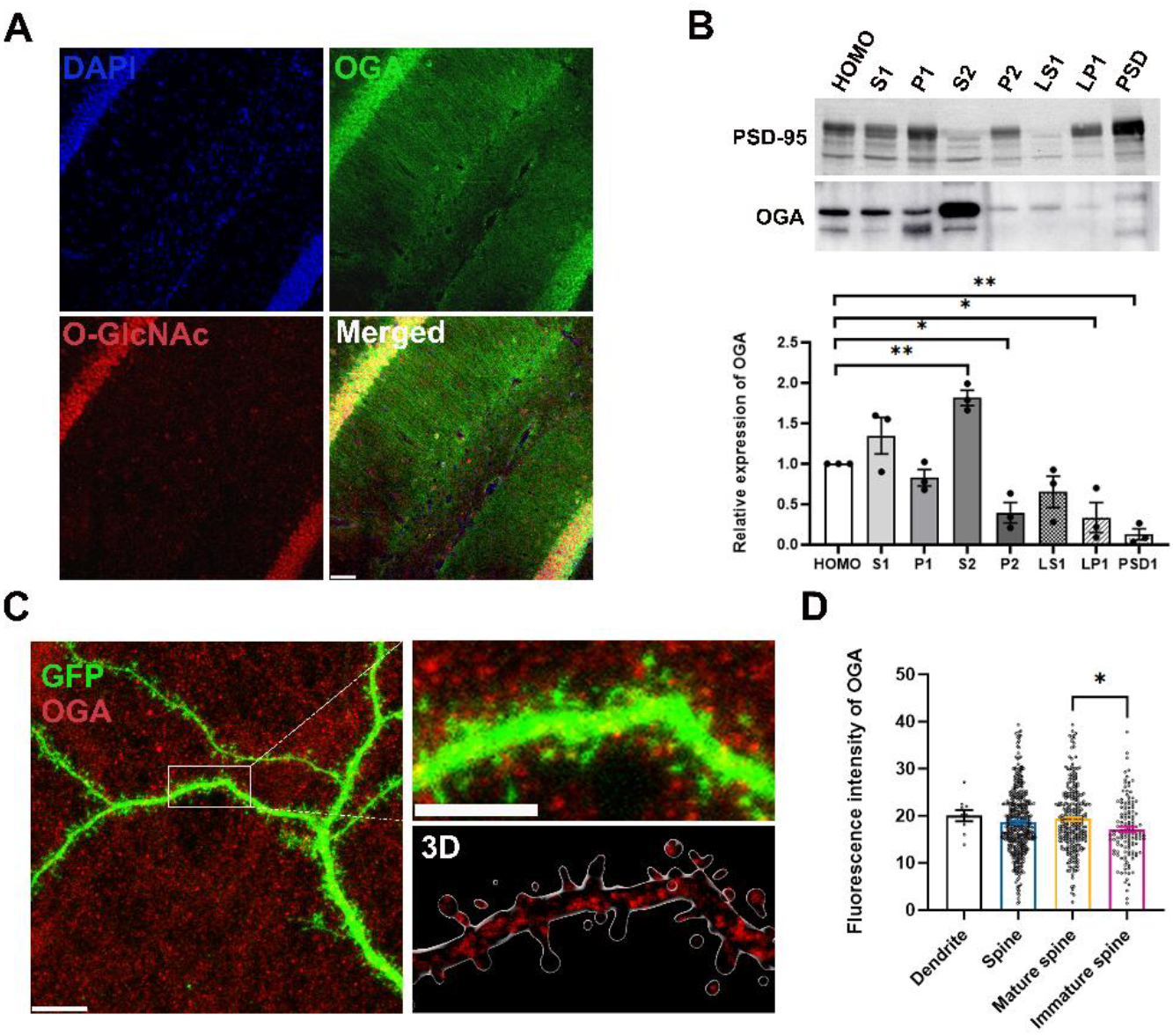
OGA is present in dendritic spines. A, Immunofluorescence image of the hippocampus. OGA (green) O-GlcNAc (red) and DAPI (blue). Scale bars 50 μm. B, Western blot of PSD fractions from brain. Quantification of expression of OGA in fractions. Data are presented as means S.E.M (*P < 0.05, **P < 0.01, ***P < 0.001.; one-way ANOVA). C, Immunostaining of the subcellular localization of OGA (red) in cultured cortical neurons (at DIV14). GFP stained the overall morphology of neurons. The scale bars represent 10 μm in (C) and 5 μm in higher magnification image and 3D image. D, Quantification of expression (C) of OGA in spines Data are presented as means S.E.M (*P < 0.05, **P < 0.01, ***P < 0.001.; one-way ANOVA). Data are presented as means S.E.M.

### Overexpression of OGA affects gross neuronal morphology

The shape, structure, and connectivity of nerve cells are critical determinants of neuronal function [28]. To examine whether OGA affects neuronal morphology at a later developmental stage, we sparsely overexpressed OGA in hippocampal cultures by transfecting neurons at DIV14 with either a GFP-expressing plasmid and an empty plasmid (WT) or a plasmid encoding OGA. Immunocytochemistry was performed to label OGA and O-GlcNAc using specific antibodies, while neurons were identified based on positive GFP staining (Fig. 2A). We analyzed OGA and O-GlcNAc intensity specifically within the soma. Quantitative analysis revealed that OGA overexpression led to a reduction in O-GlcNAc expression levels (Fig. 2B), confirming the efficacy of our manipulation. To assess the impact of OGA on neuronal morphology, we conducted Sholl analysis. Interestingly, OGA overexpression altered aspects of neuronal branching (Fig. 2A, C), the systematic analysis revealed that overexpression of OGA resulted in decreased dendritic branching compared to the control group. Collectively, these findings indicate that while OGA overexpression reduces O-GlcNAc levels, it impacts overall neuronal morphology.

**Figure 2.**
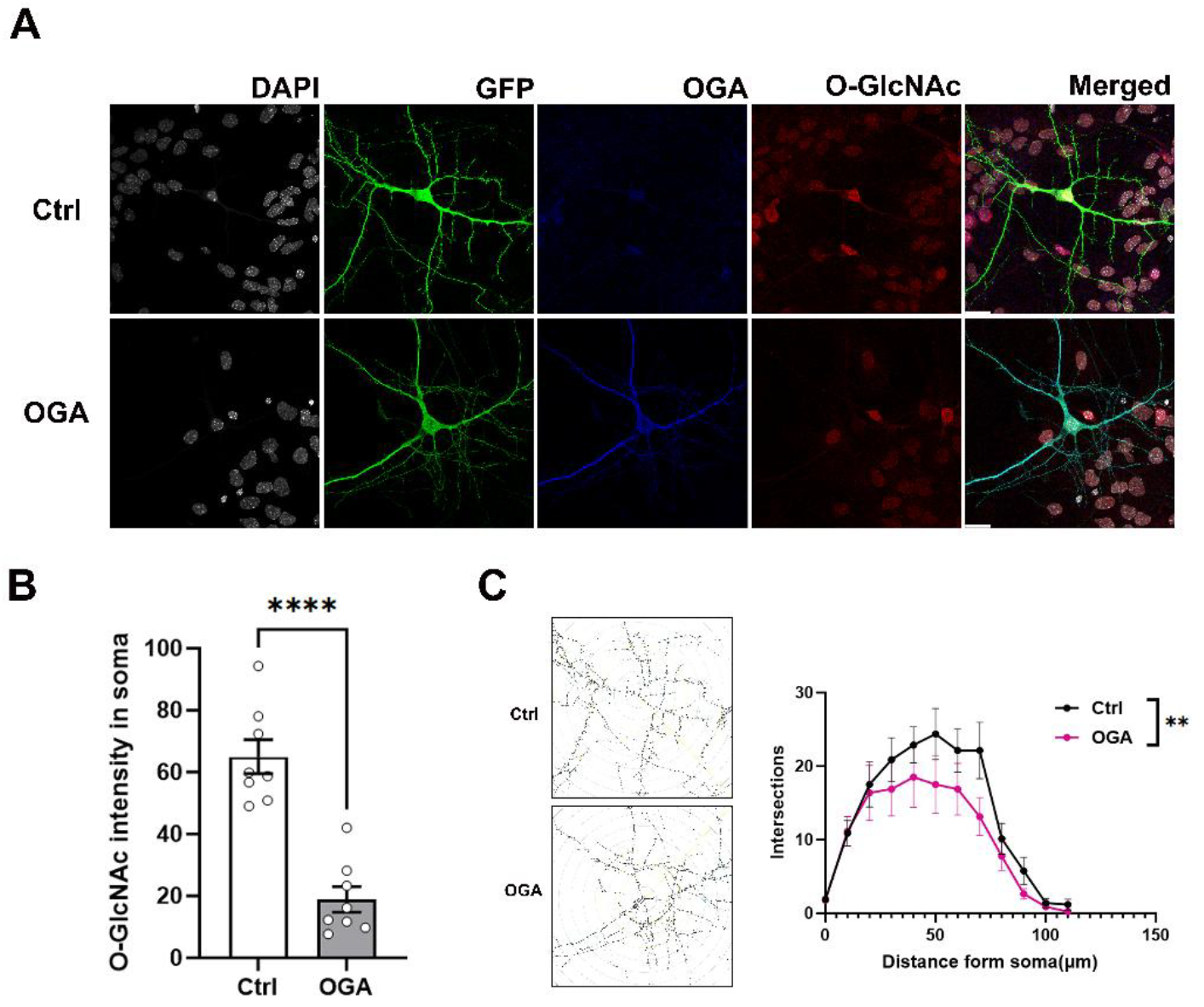
Overexpression of OGA affects gross neuronal morphology. A, Immunocytochemistry of GFP expressing hippocampal neurons stained with antibodies against OGA (blue) and O-GlcNAc (red) at DIV14. Scale bars 20 μm. B, Quantification of O-GlcNAc in hippocampal neurons(soma). (Ctrl n = 8, OGA n = 8). Data are presented as means S.E.M (*P < 0.05, **P < 0.01, ***P < 0.001.; unpaired Student’s t test). C, Sholl dendritic analysis of reconstructed neurons was performed by placing a series of concentric circles spaced at 10 μm intervals and centered on the soma. The total number of intersections in different groups was analyzed. Data was obtained through paired t-test analysis. **P < 0.01, (n = 8).

### OGA promotes mature spines in developing neurons

The expression and synaptic localization of OGA suggest its potential role in dendritic spine morphogenesis. To investigate whether OGA overexpression affects dendritic spine development, we transfected hippocampal neurons at DIV7—an early, immature stage when spine formation is emerging—with plasmids encoding OGA and GFP. GFP expression provided a general outline of neuronal morphology (Fig. 3A). OGA overexpression led to a significant increase in spine density compared to the control group (Fig. 3C). Dendritic spine morphology is typically classified based on size and shape into two main categories—mature spines characterized by mushroom spines with a wide spine head and immature spines that are uniformly thin—which reflect differences in maturity, structural dynamics, and functional properties[29] (Fig. 3B).

**Figure 3.**
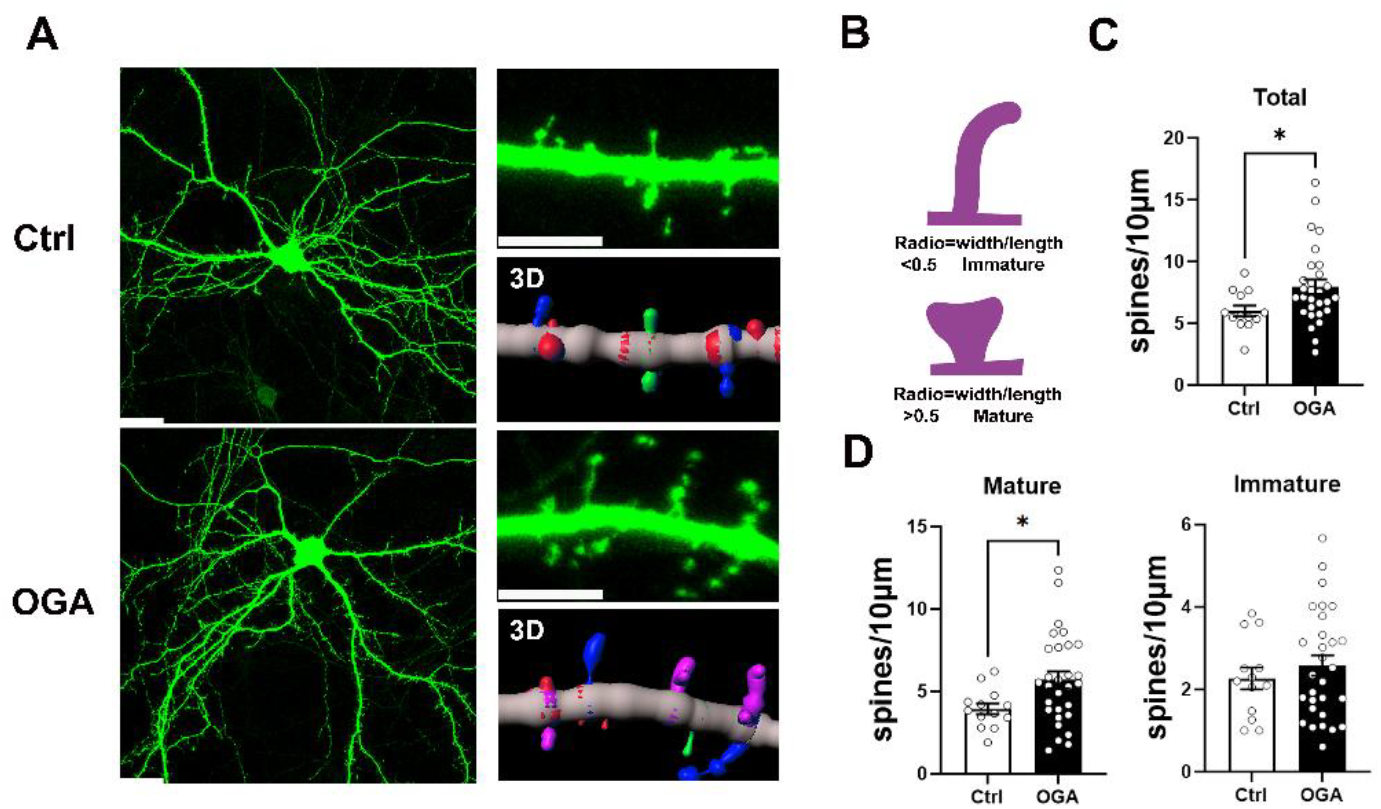
OGA promotes mature spine in developing neurons. A, Expression of GFP (green) in dendrites from WT and OGA overexpressing hippocampal neurons. Right: higher magnification image and corresponding 3D reconstruction. The scale bars represent 20 μm in (A) and 5 μm in higher magnification image. B, Schematic of measurements taken to quantify spine shape. C, Quantification of spine density (Ctrl n = 13, OGA n = 29). D, Quantification of the proportion of mature and immature spines along dendrites. (Ctrl n = 13, OGA n = 29). Data are presented as means S.E.M (*P < 0.05, **P < 0.01, ***P < 0.001.; unpaired Student’s t test).

Overexpression of OGA significantly increased the density of mature spines on secondary dendrites, while the density of immature spines remained unchanged (Fig. 3D). In summary, these findings suggest that OGA overexpression promotes dendritic spine maturation and increases overall spine density during early neuronal development.

### OGA decreases synapse size in developing neurons

Spine number and morphology are associated with synapse number [5, 30]. The formation of synaptic connections between growing axons and their appropriate targets is essential for the proper functioning of the nervous system[31-33]. To further investigate the role of OGA, we expressed GFP and OGA in cultured hippocampal neurons (Fig. 4A), prompting us to label cells with antibodies against the excitatory presynaptic marker vGluT1 and the postsynaptic marker PSD-95 (Fig. 4A).

**Figure 4.**
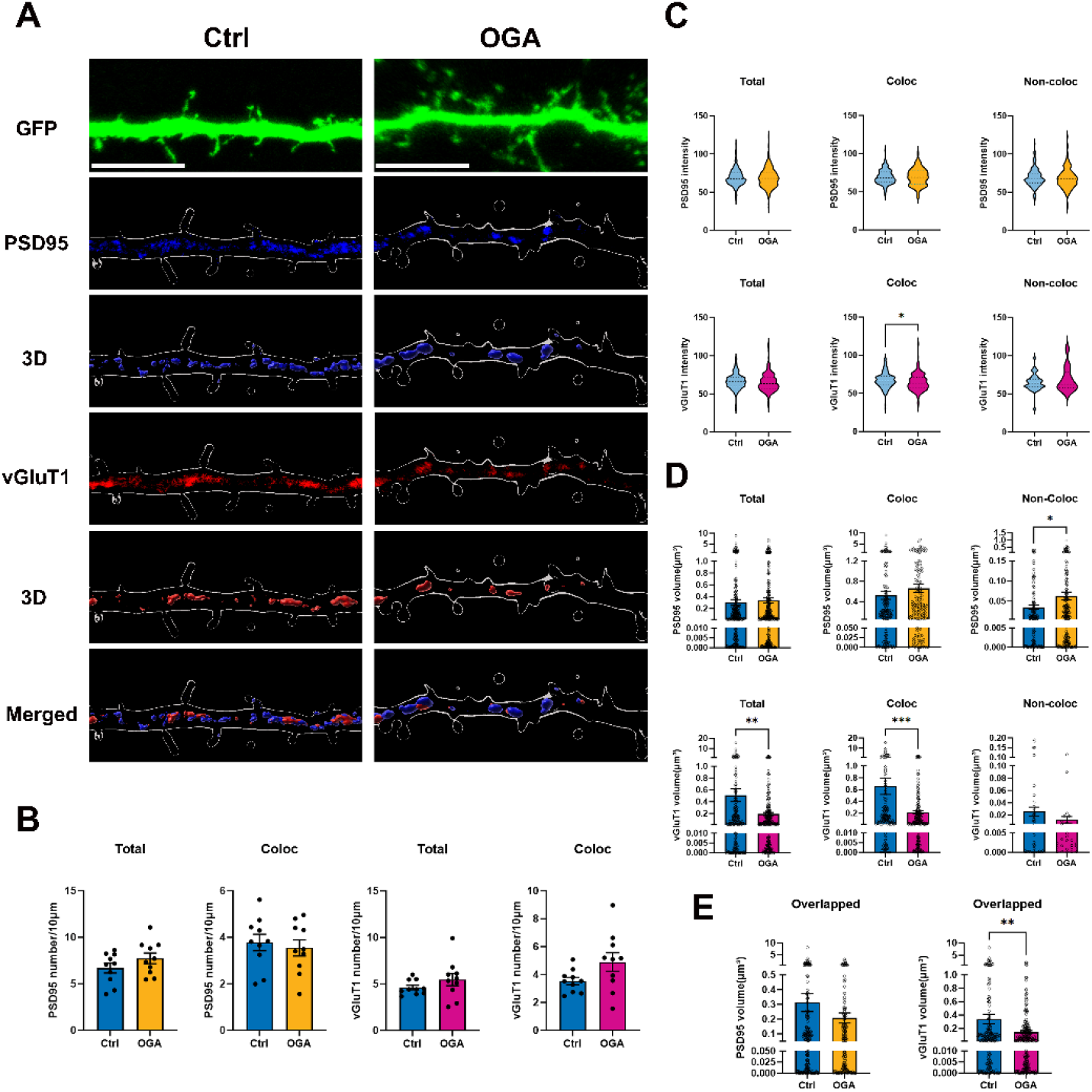
OGA decreases synapse size in developing neurons. A, Representative images of cultured hippocampal neurons transfected at DIV7 with expressing GFP alone (Control) or co expressing GFP with OGA (OGA overexpression). Neurons were analyzed by triple immunofluorescence labeling for GFP (green), PSD95 (blue), and vGluT1 (red). Scale bar, 5μm (applies to all images). B-E, Quantification graphs of the effects of OGA overexpression in neurons on density (B), intensity (C), and size (D, E) of PSD-95 or vGluT1 puncta. Data are presented as S.E.M (*P < 0.05, **P < 0.01, ***P < 0.001.; unpaired Student’s t test).

Quantitative analysis revealed no significant differences in the number, size, or intensity of PSD-95 puncta associated with vGlut1 between OGA-overexpressing and control neurons (Fig. 4B-E). However, the size of PSD-95 puncta lacking colocalization with vGluT1 was significantly increased in OGA-overexpressing neurons compared to controls (Fig. 4B). Additionally, vGluT1 labeling was used to further identify excitatory synapses. While the number of vGluT1 puncta remained comparable between OGA-overexpressing and wild-type neurons (Fig. 4B), their size was significantly reduced, indicating a decrease in presynaptic terminal size (Fig. 4D, E). Moreover, the intensity of vGluT1 puncta colocalized with PSD-95 at synapses was also diminished in OGA-overexpressing neurons (Fig. 4C). Taken together, these findings suggest that increased OGA expression leads to a reduction in excitatory synapse size, primarily by affecting presynaptic structural properties.

### OGA promotes mature spines in mature neurons

To further investigate the role of OGA in dendritic spine morphogenesis, we examined its effects not only in developing neurons (DIV7) but also in more mature cultured hippocampal neurons at DIV14(Fig. 5A). Consistent with our observations at DIV7, OGA overexpression at DIV14 resulted in a significant increase in spine density on secondary dendrites (Fig. 5B). Furthermore, spine classification analysis revealed a significant increase in the density of mature and immature spines compared to the control group (Fig. 5C). These findings suggest that OGA has a pronounced effect on mature and immature spines in developed neurons, promoting their formation and stability.

**Figure 5.**
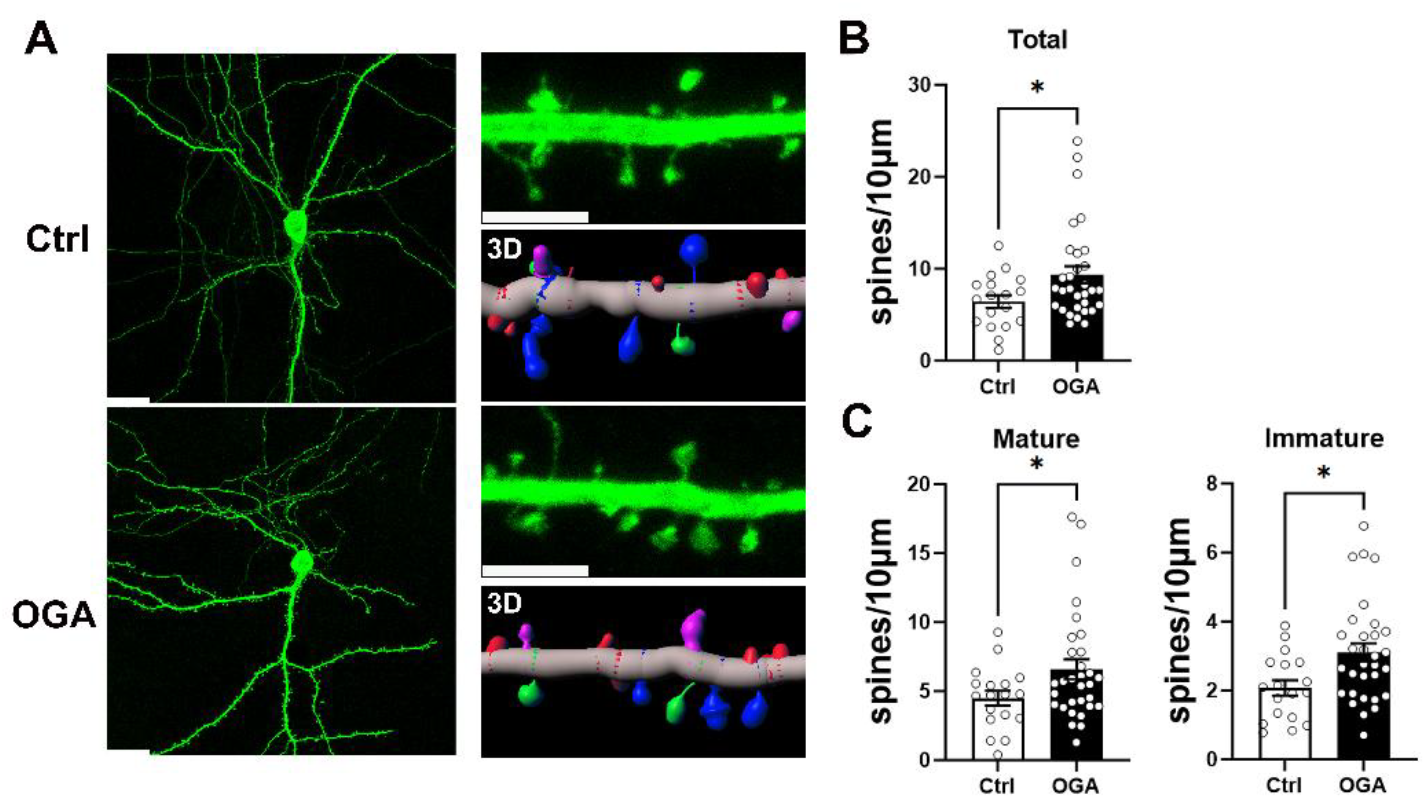
OGA promotes mature spines in mature neurons. A, Expression of GFP in dendrites from WT and OGA overexpressing hippocampal neurons. Right: higher magnification image and corresponding 3D reconstruction. The scale bars represent 20 μm in (A) and 5 μm in higher magnification images. B, Quantification of spine density (Ctrl n = 18, OGA n = 31). C, Quantification of the number of mature and immature spines along dendrites. (Ctrl n = 18, OGA n = 31). Data are presented as means S.E.M (*P < 0.05, **P < 0.01, ***P < 0.001.; unpaired Student’s t test).

### OGA decreases synapse size in mature neurons

We also immunolabeled hippocampal neurons with antibodies against presynaptic and postsynaptic markers and analyzed synapse density and size in mature neurons (DIV14) (Fig. 6A). Quantitative analysis revealed no significant differences in the number of PSD-95 or vGluT1 puncta between OGA-overexpressing and control neurons (Fig. 6B). However, the size of PSD-95 puncta (postsynaptic) and vGluT1 puncta (presynaptic) was significantly reduced, both in total and at colocalized synapses (Fig. 6D). Furthermore, the overlapping volume of PSD-95 and vGluT1 puncta was significantly decreased (Fig. 6E), suggesting that OGA overexpression leads to a reduction in synapse size in mature neurons. Additionally, fluorescence intensity measurements showed a significant decrease in both PSD-95 and vGluT1 puncta in OGA-overexpressing neurons compared to controls (Fig. 6C). In summary, these findings demonstrate that OGA plays a critical role in regulating synapse size in mature neurons, further supporting its involvement in synaptic remodeling.

**Figure 6.**
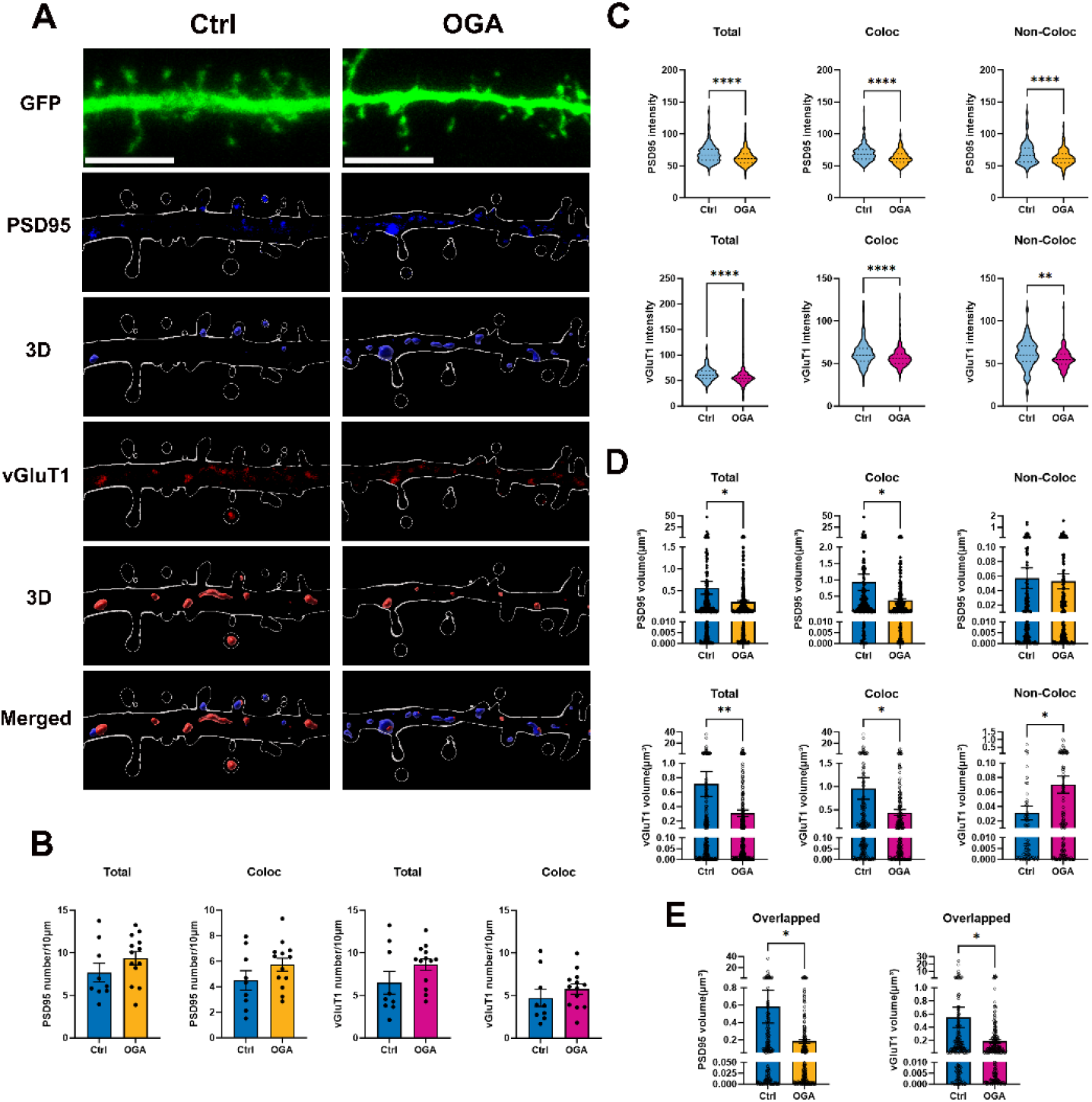
OGA decreases synapse size in mature neurons. A, Representative images of cultured hippocampal neurons transfected at DIV14 with expressing GFP (green) alone (Control) or co expressing GFP with OGA (OGA overexpression). Neurons were analyzed by triple immunofluorescence labeling for GFP (green), PSD95 (blue), and vGluT1 (red). Scale bar, 5μm (applies to all images). B-E, Quantification graphs of the effects of OGA overexpression in neurons on density (B), intensity (C), and size (D, E) of PSD-95 or vGluT1 puncta. Data are presented as means S.E.M (*P < 0.05, **P < 0.01, ***P < 0.001.; unpaired Student’s t test).

### OGA regulates AMPAR subunit composition and synaptic expression

Spine and synapse plasticity depends on AMPAR[34, 35]. AMPARs mediate the majority of fast excitatory synaptic transmission in the brain, and their precise trafficking is essential for excitatory neurotransmission, synaptic plasticity, and the formation and modification of neural circuits during learning and memory [36-38]. AMPARs are tetrameric complexes composed of GluA1–4 subunits. While the most abundant forms are GluA1/GluA2 and GluA2/GluA3 heteromers, GluA2-lacking receptors play a unique role in synaptic plasticity [39, 40]. They are permeable to calcium and have been associated with synapse and spine stabilization[41, 42]. To investigate the role of OGA in AMPAR regulation, we overexpressed OGA in cultured hippocampal neurons by transfecting plasmids expressing either GFP alone or GFP together with OGA at DIV7 or DIV14 (Fig. 7A, C). Spine classification allowed us to quantify receptor subunit localization in total, mature, and immature spines. At DIV7, OGA overexpression significantly increased GluA1 levels, while GluA2 and GluA3 levels were reduced in dendritic spines (Fig. 7B). In mature neurons (DIV14), GluA2 expression remained significantly decreased compared to controls in mature and immature spines. Notably, GluA1 expression was reduced, whereas GluA3 levels were elevated in OGA-overexpressing neurons (Fig. 7D), suggesting a differential effect of OGA on AMPAR subunit composition during neuronal maturation, with consistent downregulation of GluA2 in both developing and mature neurons.

**Fig. 7.**
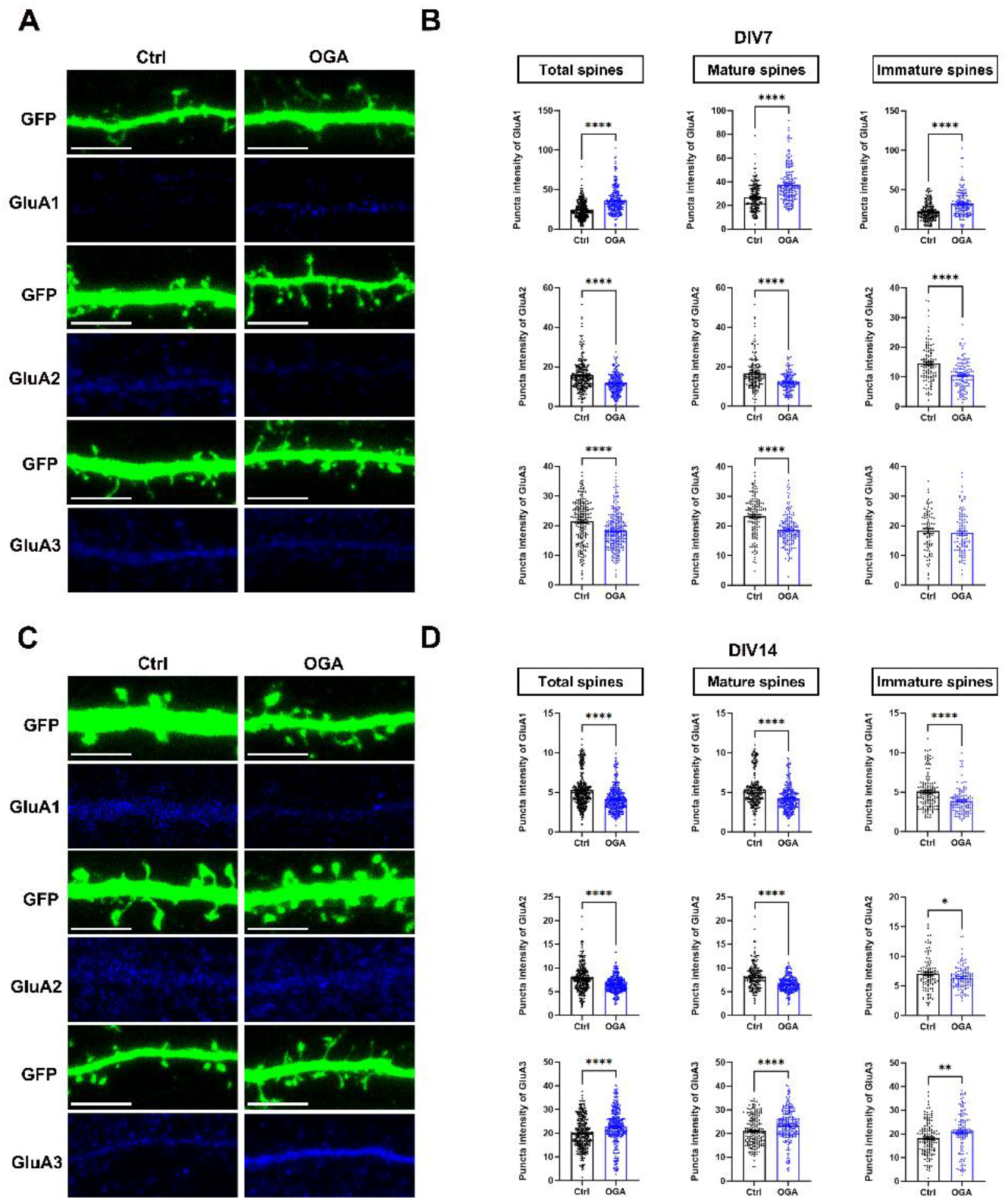
OGA regulates AMPAR subunit composition and synaptic expression. A, Immunohistochemistry images of GFP expressions in WT and OGA overexpressing hippocampal neurons transfected at DIV7. Neurons were analyzed by immunofluorescence labeling for GluA1 (blue), GluA2 (blue) or GluA3(blue). Scale bar, 5μm (applies to all images). B, Quantification of the total expression of GluA1(upper), GluA2(middle) and GluA3(lower) in the spines of individual hippocampal neurons. C Immunohistochemistry images of GFP expression in WT and OGA overexpressing hippocampal neurons transfected at DIV14. Neurons were analyzed by immunofluorescence labeling for GluA1 (blue), GluA2 (blue) or GluA3(blue). Scale bar, 5μm (applies to all images). D, Quantification of the total expression of GluA1(upper), GluA2(middle) and GluA3(lower) in the spines of individual hippocampal neurons. Scale bar, 5μm (applies to all images). Data are presented as means S.E.M (*P < 0.05, **P < 0.01, ***P < 0.001.; unpaired Student’s t test).

Together, these data suggest a model in which OGA neurons exhibit altered neuronal morphology, an increased number of dendritic spines, reduced synapse size, and a selective loss of GluA2-containing AMPARs (Fig. 8). Our results suggest that OGA regulates excitatory synaptic structure and receptor composition, potentially influencing synaptic strength and plasticity.

**Fig. 8.**
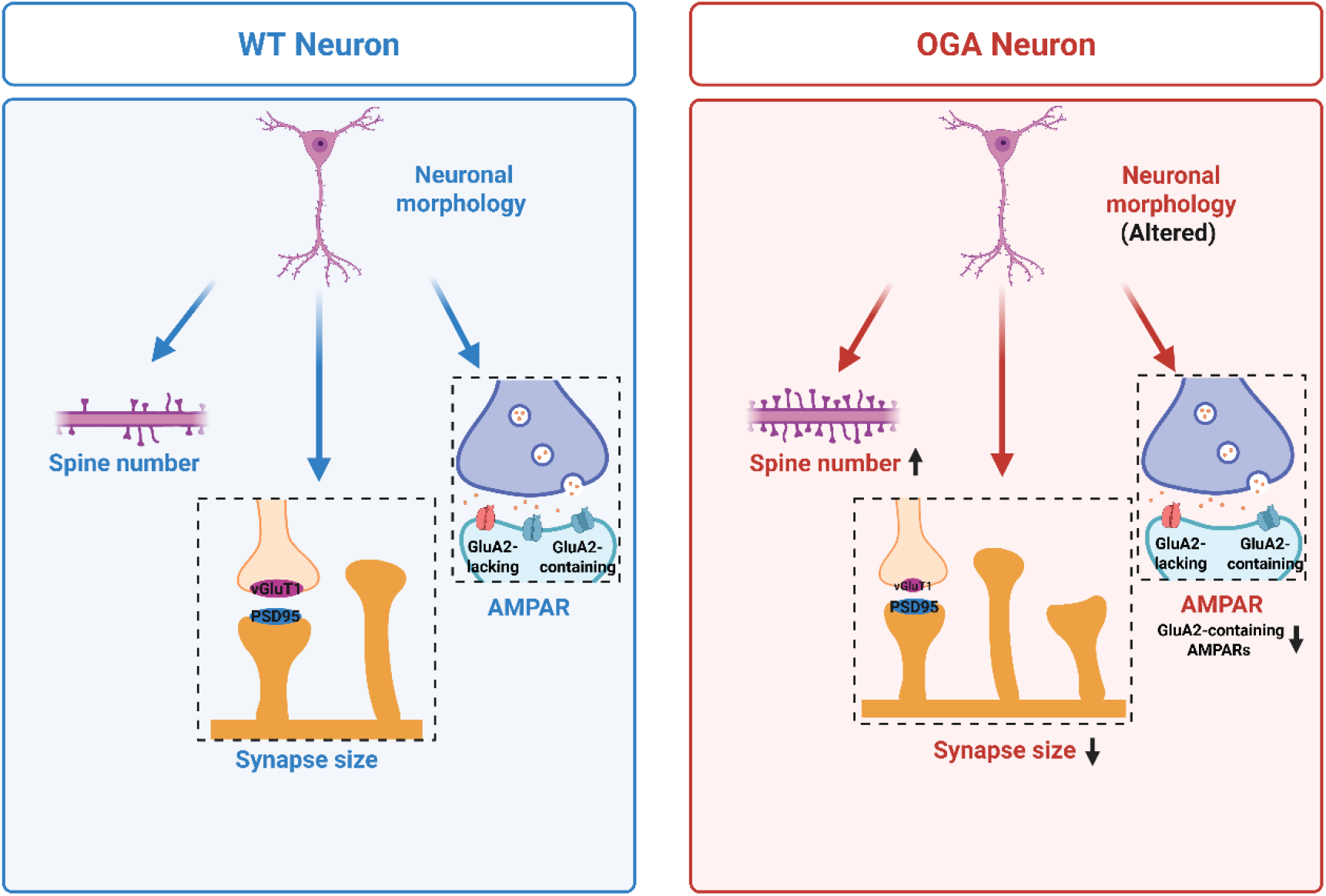
A model of altered Synaptic Properties in OGA Neurons. Schematic model illustrates synaptic differences between wild-type (WT) and O-GlcNAcase (OGA) neurons. In WT neurons (left, blue panel), neuronal morphology, dendritic spine number, synapse size, and AMPA receptor (AMPAR) composition are maintained under normal conditions. In OGA neurons (right, red panel), altered neuronal morphology is accompanied by an increased spine number, reduced synapse size, and decreased GluA2-containing AMPARs at synapses.

## Discussion

Our study provides new insights into the role of O-GlcNAcase (OGA) in dendritic spine morphogenesis and synaptic function. Through a series of biochemical and imaging analyses, we demonstrate that OGA is localized in dendritic spines and plays a crucial role in regulating spine maturation and synapse size. Our findings suggest that OGA modulates synaptic plasticity by influencing the structural and molecular properties of excitatory synapses, particularly through its effects on AMPA receptor subunit composition.

We first established that OGA is predominantly localized in the soma and dendrites of neurons and is present within dendritic spines, while being largely excluded from the postsynaptic density (Fig. 1). Overexpression of OGA altered neuronal morphology and significantly increased the density of mature spines in both developing and mature neurons (Figs. 2, 3, and 5). Surprisingly, this morphological change was accompanied by a reduction in synapse size, as indicated by decreased PSD-95 and vGluT1 puncta size and fluorescence intensity (Figs. 4 and 6). Furthermore, OGA consistently downregulated GluA2-containing AMPARs across both developmental stages, while GluA1 and GluA3 levels were differentially affected (Fig. 7).

The observed increase in mature spine density suggests that OGA contributes to postsynaptic structural stabilization. The concurrent downregulation of GluA2, a calcium-impermeable AMPAR subunit essential for synaptic stability, indicates a shift toward GluA2-lacking, calcium-permeable AMPARs. This shift likely increases calcium influx at synapses, activating signaling pathways involved in actin cytoskeleton remodeling, a key process in spine growth and maturation[41, 42]. Thus, OGA may regulate spine and synapse development by modulating AMPAR subunit composition and calcium dynamics. The shift in AMPAR composition may explain how OGA increases spine maturity while inhibiting synapse maturity. GluA2-containing AMPA receptors promote synapse maturation through their extracellular domain[43, 44]. In contrast, GluA2-lacking receptors have been shown to affect early phases of spine morphogenesis associated with long term potentiation of excitatory synapses[45-47]. These results are strengthened by previous electrophysiological studies in brain slices using drugs to inhibit OGA which indicated that OGA may regulate synaptic plasticity through affecting the ratio between GluA2-lacking and GluA2-containing receptors in synapses[15, 48]. Our results present molecular and structural evidence for how OGA affects synaptic plasticity. We also show evidence indicating that the molecular mechanism by which OGA affects synaptic plasticity depends on neuronal development. While changes in AMPAR composition provide a plausible mechanism, OGA may also influence other proteins involved in synapse formation and maintenance. Given that O-GlcNAc is involved in cellular signaling and protein regulation[49], OGA may directly affect scaffolding proteins, adhesion molecules, or actin-regulating factors that shape synaptic architecture.

Some of our findings on the effects of overexpression OGA – functionally reducing general O-GlcNAc levels – are similar to the effects of decreasing general O-GlcNAc levels by knocking out O-GlcNAc transferase (OGT), the enzyme responsible for adding O-GlcNAc, which has been shown to regulate synaptic protein stability and neurotransmission[14]. Reducing total O-GlcNAc by either OGT deletion or OGA overexpression inhibits synaptic development and promotes GluA2-lacking while removes GluA2-containing receptors. However, OGA overexpression increased spine maturation when OGT deficiency decreases spine maturation[14]. These findings support the broader concept that balanced O-GlcNAc cycling is critical for maintaining synaptic structure and function. They also show that a mechanistic understanding of the function of O-GlcNAc cycling requires identification of what O-GlcNAc sites mediates the effect of OGT and OGT on synaptic plasticity.

Importantly, our data aligns with in vivo observations in OGA heterozygous knockout mice, which reported alterations in spine density and synaptic organization[50], reinforcing the physiological relevance of OGA in synaptic plasticity. In addition to identifying mechanisms, our use of an acute overexpression model in cultured hippocampal neurons provides a dynamic perspective that complements chronic genetic manipulation approaches. This temporal specificity reveals how rapid shifts in O-GlcNAc levels can influence synaptic development and receptor composition in real time. Our data show that OGA is an important mediator of several forms of synaptic plasticity not only during development but also in mature neurons.

Understanding the role of OGA in synaptic function has important implications for cognitive health and neurological disorders. Dendritic spine abnormalities are associated with conditions such as intellectual disability, Alzheimer’s disease, autism spectrum disorders, and schizophrenia[51, 52]. They all involve disruptions in synaptic connectivity and plasticity. Emerging evidence links OGA to these disorders through both genetic and functional studies. Distinct OGA splice variants in the human brain may differentially impact neuronal signaling and development, and mutations in OGA have been associated with intellectual disability[53], underscoring the importance of OGA regulation for cognitive health. Together with our data, these findings support the model that OGA directly influences synaptic structure and function. Targeting OGA—whether through modulation of expression, enzymatic activity, or splice variant usage—may therefore offer novel therapeutic opportunities for treating neurodevelopmental and neurodegenerative disorders.

In summary, our study identifies OGA as a key regulator of dendritic spine maturation and synaptic remodeling. By modulating spine density, synapse size, and AMPAR composition, OGA contributes to the fine-tuning of excitatory synaptic connections. These findings advance our understanding of O-GlcNAc-mediated synaptic regulation and highlight OGA as a promising target for therapeutic intervention in neurological disorders characterized by impaired synaptic plasticity.

## Methods

### Animals

We used C57BL6 mice. Animals were housed under a normal 12 h light–dark cycle with ad libitum water and food. The ambient temperature was 25 °C and humidity was 50%. Both male and female mice were used in the study. We have complied with all relevant ethical regulations for animal use. All experimental procedures were approved and conducted in accordance with the regulations of the Local Animal Ethics Committee at Umea University.

### Primary neuronal cultures

Mouse primary hippocampal neurons were prepared from E18 mice brains, as described previously[54, 55] [56], Neurons were plated on coverslips coated with poly-L-lysine at a density of 300,000 cells/well and grown in NM5 [neurobasal growth medium (Gibco) with 5% (vol/vol) serum, 2% (vol/vol) B27 (Invitrogen), 2% (vol/vol) B27 (Invitrogen)]. Then, 2 h after plating, the medium was replaced with NM0[neurobasal growth medium (Gibco) with 5% (vol/vol) serum, 2% (vol/vol) B27 (Invitrogen), 2% (vol/vol) B27 (Invitrogen)]. Cultures were maintained in an incubator at 37 °C under 5% CO2/95% air and 90% humidity.

### Transfection

At DIV7 or DIV14, primary hippocampal neurons in 12-well plates were transfected with Lipofectamine 2000 (2 µl/well; Thermo Fisher Scientific, 10696343) and 1.5 µg plasmid DNA/well in Neurobasal medium (NM0 + B27, no serum/antibiotics). DNA and Lipofectamine were each diluted to 50 µl/well in plain Neurobasal, incubated separately for 5 min, then combined (1:1) and incubated for 20 min at 37 °C. Complexes (100 µl/well) were applied for 45 min, after which wells were rinsed with plain Neurobasal and returned to conditioned medium.

### Western blot

Homogenized brain tissues were harvested in RIPA buffer (50mM Tris–HCl, pH 8.0, 150 mM NaCl, 1% NP-40, 1% deoxycholate, 0.1% SDS, 1mM EDTA) supplemented with a protease inhibitor Phosphatase and O-GlcNAcases inhibitor. The resulting lysates were centrifuged at 4°C at 13000 rpm, and the supernatants were used for further analysis. Protein samples were mixed in the loading buffer and denatured by boiling for 10 min at 90°C. Western blotting was performed according to standard procedures. Briefly, the protein samples were subjected to SDS-PAGE electrophoresis and transferred into polyvinylidene fluoride (PVDF) membranes, membranes were blocked in 3% bovine serum albumin (BSA). The membranes were incubated with the following antibodies: OGA (Thermo Fisher Scientific, 16813494, 1:1000), anti-PSD95(NeuroMab, K28/43, 1:1000) overnight at 4 °C and then with secondary HRP-conjugated antibodies (anti-rabbit (Thermo Fisher Scientific, 31460, 1:5,000) or anti-mouse (Thermo Fisher Scientific, 31430, 1:5,000)) for 1 h at room temperature. The protein signals were visualized with the western blotting detection reagent (Thermo Fisher Scientific, 34580) and scanned with the Sapphire biomolecular imager (Azure Biosystems, IS4000).

### Immunochemistry

Brain sections (50 µm; 4 per well) were processed in 12-well plates using 0.5 mL solution per well for incubations and 1 mL for washes. Sections were blocked and permeabilized overnight at 4 °C in PBS containing 5% normal goat serum and 0.25% Triton X-100 with gentle rocking, then incubated with primary antibodies overnight at 4 °C in the same buffer. After three 45 min washes in PBS/Triton X-100 (0.25%), sections were incubated with secondary antibodies for 2 h at room temperature or overnight at 4 °C, with DAPI (Sigma, D9542-10MG; 1:1000) added to the secondary solution. Following three additional washes, sections were mounted in Fluoromount-G (Thermo Fisher Scientific, 15586276), dried overnight in darkness, and stored at 4 °C or −20 °C.

Cultured neurons were fixed with 4% paraformaldehyde, 4% sucrose, for 10-15 min at 4°C. Then neurons were permeabilized with 0,25% Tx-100 for 10 min at 4°C and blocked in blocking buffer (Normal goat serum, Vector Laboratories, S-1000-20) for 1 h at room temperature. This was followed by incubation with primary antibodies for 2h at room temp. After washing, secondary antibodies were applied for 1 h at room temperature. After washing for 3 × 10 min in PBS, samples were mounted in fluorescence mounting medium (Fluoromount-G), and stored at 4 °C.

Primary antibodies for immunocytochemistry: anti-O-GlcNAc (Thermo Fisher Scientific, MA1-072, 1: 500). Anti-OGA (Thermo Fisher Scientific, 16813494, 1:1000), anti-GFP (Aves labs, GFP-1020, 1:1000), anti-PSD95(NeuroMab, K28/43, 1:1000), anti-vGluT1(Millipore, AB5905, 1:1000), anti-GluA1(NeuroMab, N355/1, 1:100), anti-GluA2(NeuroMab, L21/32, 1:100), anti-GluA3(Thermo Fisher Scientific, 32-0400, 1:1000), anti-Chicken secondary antibody(Azure Biosystems, AC2209, 1:2000), anti-Guinea Pig secondary antibody(Thermo Fisher Scientific, A21450, 1:2000), anti-Mouse secondary antibody(Thermo Fisher Scientific, A21422, 1:2000), anti-Rabbit secondary antibody(Thermo Fisher Scientific, A21428, 1:2000).

### Image acquisition and analysis

Confocal images of the neuronal cell cultures and brain sections were captured using a Leica SP8 FALCON Confocal. z-stack confocal images covering ∼50µm thickness were acquired at 0.5-µm intervals. The original images were analyzed using Imaris 10.2.1 (Bitplane). FilamentTracer in Imaris was used to trace and analyze the spine density and maturation. The number and intensity of the protein marker-positive puncta were quantified. PSD95 and vGluT1 within the GFP^+^ neuron were 3D-reconstructed and then assessed automatically by Imaris software.

To better present our results, moderate linear adjustments were applied to the entire image (rather than to specific areas). For all images displayed, contrast and brightness were applied linearly across the entire image. All images (Control and OGA groups) were captured under the same settings and processed identically using Leica X Office software.

### Statistics

GraphPad Prism 10 (GraphPad Software) was used for statistical analyses. Comparisons between two groups were performed using Student’s T-tests. Data was presented as individual data points and as the mean value ± SEM. Significance levels were set to: ***p < 0.0001; ***p < 0.001; **p < 0.01; *p < 0.05.

